# RIG-I and PKR, but not stress granules, mediate the pro-inflammatory response to Yellow fever virus

**DOI:** 10.1101/2020.02.05.935486

**Authors:** Guillaume Beauclair, Felix Streicher, Daniela Bruni, Ségolène Gracias, Salomé Bourgeau, Laura Sinigaglia, Takashi Fujita, Eliane F. Meurs, Frédéric Tangy, Nolwenn Jouvenet

## Abstract

Yellow fever virus (YFV) is an RNA virus primarily targeting the liver. Severe YF cases are responsible for hemorrhagic fever, plausibly precipitated by excessive pro-inflammatory cytokine response. Pathogen recognition receptors (PRRs), such as the cytoplasmic RIG-I-like receptors (RLRs), and the viral RNA sensor PKR are known to initiate a pro-inflammatory response upon recognition of viral genomes. Here, we sought to reveal the main determinants responsible for the acute cytokine expression occurring in human hepatocytes following YFV infection. Using a RIG-I-defective human hepatoma cell line, we found that RIG-I largely contributes to cytokine secretion upon YFV infection. In infected RIG-I-proficient hepatoma cells, RIG-I was localized in stress granules. These granules are large aggregates of stalled translation preinitiation complexes known to concentrate RLRs and PKR, and are so far recognized as hubs orchestrating RNA virus sensing. Using PKR-deficient hepatoma cells, we found that PKR contributes to both stress granule formation and cytokine induction upon YFV infection. However, stress granules disruption did not affect the cytokine response to YFV infection, as assessed by siRNA-knockdown-mediated inhibition of stress granule assembly. Finally, no viral RNA was detected in stress granules using a fluorescence *in situ* hybridization approach coupled with immunofluorescence. Our findings suggest that both RIG-I and PKR mediate pro-inflammatory cytokine induction in YFV-infected hepatocytes, in a stress granule-independent manner. Therefore, by showing the uncoupling of the early cytokine response from the stress granules formation, our model challenges the current view by which stress granules are required for the mounting of the acute antiviral response.

**Importance:** Yellow fever is a mosquito-borne acute hemorrhagic disease caused by yellow fever virus (YFV). The mechanisms responsible for its pathogenesis remain largely unknown, although increased inflammation has been linked to worsened outcome. YFV targets the liver, where it primarily infects hepatocytes. We found that two RNA-sensing proteins, RIG-I and PKR, participate in the induction of pro-inflammatory mediators in human hepatocytes infected with YFV. We show that YFV infection promotes the formation of cytoplasmic structures, termed stress granules, in a PKR-, but not RIG-I-dependent manner. Whilst stress granules were previously postulated to be essential platforms for immune activation, we found that they are not required for pro-inflammatory mediators’ production upon YFV infection. Collectively, our work uncovered molecular events triggered by the replication of YFV, which could prove instrumental in clarifying the pathogenesis of the disease, with possible repercussions on disease management.

## Introduction

Flaviviruses are a group of more than seventy enveloped RNA viruses that can cause serious diseases in humans and animals (1, 2). They have provided some of the most important examples of emerging or resurging diseases of global significance. Most flaviviruses, such as dengue virus (DENV), yellow fever virus (YFV) and Zika virus (ZIKV), are arthropod-borne viruses transmitted to vertebrate hosts by mosquitoes or ticks (1). YFV is the prototype and eponymous virus of the Flavivirus genus. It is a small (40–60 nm), enveloped virus harboring a single positive-strand RNA genome of 11 kb. The genome encodes a polyprotein that is cleaved co- and post-translationally into three structural proteins: capsid (C), membrane precursor (prM) and envelope (E), and seven non-structural (NS) proteins (NS1, NS2a, NS2b, NS3, NS4a, NS4b, and NS5). The C, prM, and E proteins are incorporated into the virions, while NS proteins are found only in infected cells (3). NS proteins coordinate RNA replication, viral assembly and modulate innate immune responses, while the structural proteins constitute the virion.

YFV is endemic in the tropical regions of sub-Saharan Africa and South America. The reference strain Asibi was isolated in 1927 in West Africa from the blood of a human patient. The vaccine strain 17D was developed empirically in the 1930s, by passaging the Asibi strain in rhesus macaques, mouse and chicken embryonic tissues (4). 17D is one of the most effective vaccines ever generated. It has been used safely and effectively on more than 600 million individuals over the past 70 years (4). However, due to poor vaccine coverage and vaccine shortages, the virus continues to cause disease in endemic areas, as illustrated by recent outbreaks in Angola and Brazil (5, 6).

Yellow fever (YF) pathogenesis is viscerotropic in humans, with viral replication in the liver critical to the establishment of the disease (3). Severe YF is responsible for multi-system organ failure and viral hemorrhagic fever resulting in up to 50% fatality. Similar to Ebola hemorrhagic fever, cytokine dysregulation is thought to result in endothelial damage, disseminated intravascular coagulation and circulatory shock observed in the terminal stage of the disease. Viral hemorrhagic fever is considered as an illness precipitated by excessive pro-inflammatory cytokine response (‘cytokine storm’) (7, 8). Almost any cell type in the body can produce pro-inflammatory cytokines in response to various stimuli, such as a viral infection (9). The pattern of cytokine expression in response to infection with pathogenic strains of YFV has been described both in humans and in rhesus macaques. Studies performed on patients during a YF outbreak in the Republic of Guinea in 2000 showed that excessive production of pro-inflammatory cytokines, such as IL6 and TNFα, correlates with high viremia and disease outcome (10). The levels of these two cytokines were statistically higher in patients with fatal hemorrhagic fever than in patients who survived. Similar findings were obtained with rhesus macaques infected with the strain Dakar-HD1279 (11). The animals died of hemorrhagic fever within five days. These studies also suggest that pro-inflammatory cytokines might be produced by injured organs, especially the liver, and not by cells circulating in the blood (11).

Upon viral infection, the cytokine response is initiated by the recognition of viral genomes and viral replication intermediates by pathogen recognition receptors (PRRs) (12), such as the ubiquitously expressed cytoplasmic RIG-I-like receptors (RLRs). Three RLRs have been identified in vertebrates: melanoma differentiation-associated gene 5 (MDA5), laboratory of genetics and physiology 2 (LGP2) and retinoic acid inducible gene 1 (RIG-I) (13). Upon binding to viral nucleic acids, MDA5 and RIG-I undergo a conformational change, which allows interaction with the adaptor protein MAVS and subsequent recruitment of signaling complexes that comprises protein kinases such as TBK1, IKKα, IKKβ and IKKε (14). These events lead to the activation and nuclear translocation of the transcription factors IRF3 and NF-κB, which stimulate the rapid expression of pro-inflammatory cytokines and type I interferons (IFNs).

RIG-I plays an important role in initiating the IFN-mediated antiviral response against DENV and ZIKV (15–18). Induction of cytokines upon infection with the vaccine strain of YFV is also mediated by RIG-I in plasmacytoid dendritic cells (19). Whether RIG-I contributes to cytokine production in cells infected with wild-type YFV strains is not known.

The subcellular localization where RIG-I interacts with its ligands is not well described. Upon infection with influenza A virus lacking its non-structural protein 1 (IAVΔNS1), Newcastle disease virus or Sendai virus (20–23), RIG-I accumulates in cytoplasmic foci. Disruption of these foci, which have been identified as stress granules, reduced the abundance of *IFNB* mRNAs in IAVΔNS1-infected cells (20). Thus, stress granules may be important for RIG-I signaling and could serve as a platform for viral RNA sensing. Moreover, other viral nucleic sensors, such as MDA5 (24), LGP2 (25) and cGAS (26), are also concentrated in stress granules. Finally, numerous viruses interfere with stress granule formation, *via*, for instance, the cleavage of one of their core components by a viral protease. Thus, accumulation of viral sensors in stress granules and viral counteraction mechanisms suggest that these granules act as hubs that coordinate foreign nucleic acid sensing (27, 28).

Stress granules are non-membranous cytoplasmic structures that concentrate translationally stalled mRNAs and proteins (29). Their assembly is triggered by sensing distinct types of stress, including viral infection, by one of four kinases, resulting in phosphorylation of the eukaryotic initiation factor 2α (eIF2α). These kinases are: the heme-regulated inhibitor (HRI), which is activated by heat shock or oxidative stress (*e.g*. by reactive oxygen species (ROS) produced upon treatment with sodium arsenite); general control non-depressible protein 2 (GCN2), which reacts to UV-induced damage or amino acid deprivation; dsRNA-sensing protein kinase R (PKR); and PKR-like endoplasmatic reticulum (ER) kinase (PERK), which is activated upon ER stress (29). Stress granule assembly does not only arrest the host translation machinery, but also heavily disturb viral translation, which is further evidence for their antiviral nature (27, 28, 30). Here, we investigated the role of RIG-I, PKR and stress granules in the induction of inflammatory cytokines upon infection of hepatoma cells with YFV.

## Results

### The Asibi strain of YFV triggers the production of pro-inflammatory cytokines in hepatoma cells through RIG-I, IRF3 and NF-κB

We examined whether the YFV reference strain Asibi could trigger the expression of pro-inflammatory cytokines in human hepatoma cells through the RIG-I signaling pathway. Huh7 and Huh7.5 cells, a clone derived from Huh7 that expresses an inactive form of the viral sensor RIG-I (31), were infected with the Asibi strain for 24 or 48 hours. The abundances of *TNFA*, *IL6* and *IFNB* mRNAs increased over time with infection in Huh7 cells, while being significantly lower in Huh7.5 cells at 48 hours post-infection (hpi) (**Fig. 1A-C**). We noticed that Huh7.5 cells infected for 48 hours produced approximately three-fold more viral RNA than Huh7 cells infected for the same amount of time (**Fig. 1D**). Albeit not statistically significant, this modest effect was reproducible (**Fig. 1D**) and suggests that absence of RIG-I signaling favors viral replication. Huh7 cells secreted around 10 to 20 pg/ml of IL6 and TNFα protein after 48 hours of infection, but not after 24 hpi (**Fig. 1E, F**). By contrast, the amount of IL6 and TNFα protein secreted by cells expressing the inactive form of RIG-I was below the detection limit of the ELISA (**Fig 1E, F**). These data suggest that Asibi infection triggers the induction and secretion of pro-inflammatory cytokines in a RIG-I-dependent manner in human hepatoma cells.

**Fig. 1:**
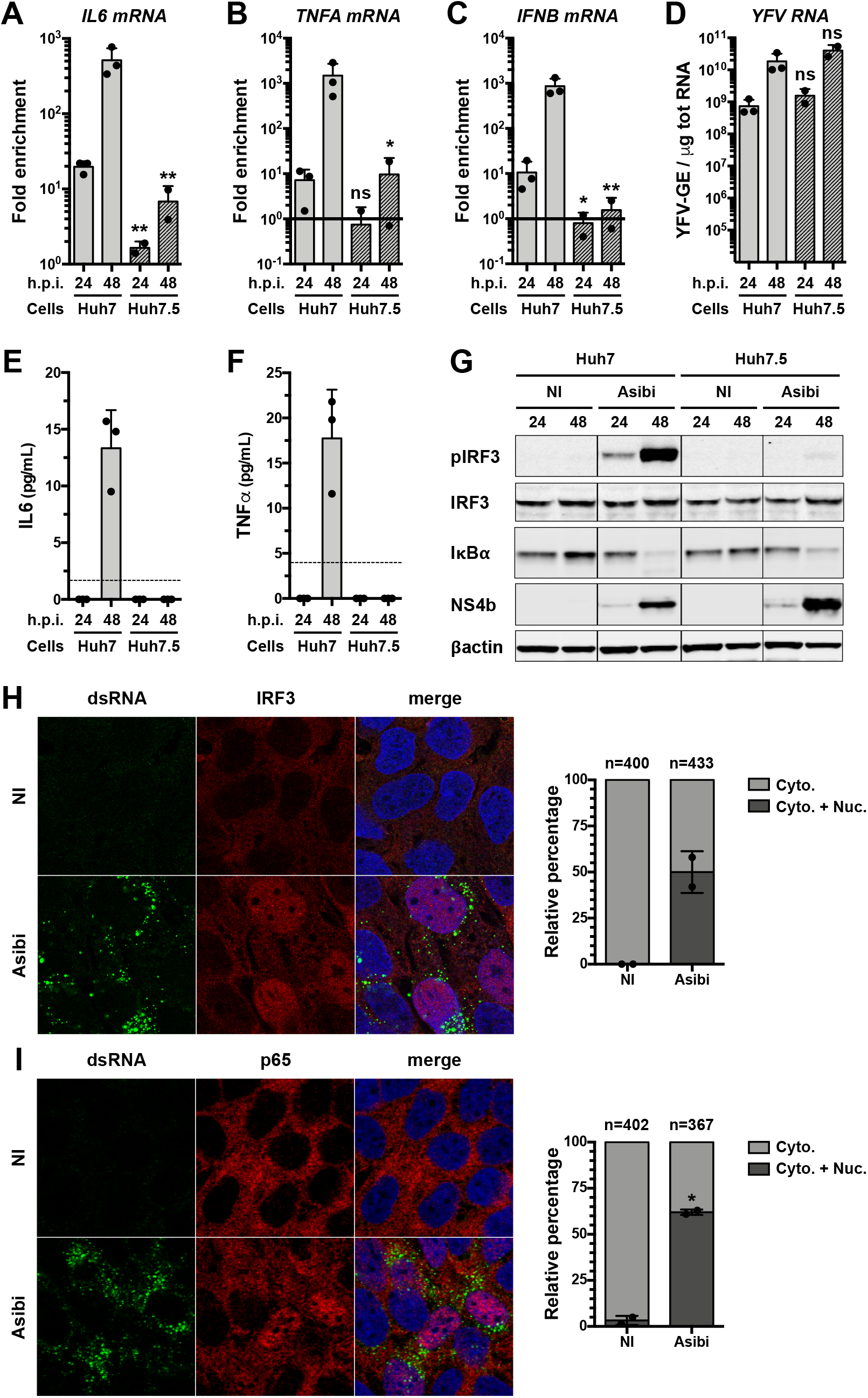
The Asibi strain of YFV triggers the production of pro-inflammatory cytokines in hepatoma cells through RIG-I, IRF3 and NF-κB. Huh7 and Huh7.5 cells were left uninfected or infected with Asibi at an MOI of 1 for 24 or 48 hours. The relative amounts of *IL6* (**A**), *TNFA* (**B**) and *IFNB* (**C**) mRNAs were determined by RT-qPCR analysis and normalized to GAPDH mRNA and non-infected samples. The relative amounts of cell-associated viral RNA were determined by RT-qPCR analysis (**D**) and expressed as genome equivalents (YFV-GE) per μg of total cellular RNA. The data are the means ± SD of two (Huh7.5) or three (Huh7) independent experiments. ns: non-significant, *: p<0.05, **: p<0.01. (**E**, **F**) Culture media of the indicated cells were analysed by ELISA to determine the amounts of secreted IL6 (**E**) and TNFα (**F**). The data are the means ± SD of two (Huh7.5) or three (Huh7). The dashed lines indicate the limit of detection of the IL6 and TNFα ELISA (2 and 4 pg/ml, respectively). (**G**) Whole-cell lysates of the indicated cells were analysed by Western blotting with antibodies against the indicated proteins. (**H**, **I**) Huh7 cells were left uninfected (NI) or infected with YFV-Asibi for 48 hours at an MOI of 20. Cells were then stained with IRF3 (**H**) or p65 (**I**) (red), dsRNA (green) antibodies and NucBlue^®^ (blue). Around 200 cells per independent experiment were scored for nuclear translocation of IRF3 or p65 (**H** and **I**, respectively). ns: non-significant, *: p<0.05.

By Western blot analysis, we found that IRF3 was activated in Huh7 cells infected with Asibi for 24 or 48 hours, but not in non-infected cells (**Fig. 1G**). The level of IRF3 phosphorylation was higher at 48 than at 24 hpi, despite the total protein abundance being similar at these two points in time (**Fig. 1G**). IRF3 was not phosphorylated in Huh7.5 cells infected with Asibi (**Fig. 1G**). IκBα is a negative regulator of the NF-κB pathway: its degradation triggers the nuclear translocation of the NF-κB subunits p65 and p50. IκBα was degraded in Huh7 cells infected for 48 hours (**Fig. 1G**). Slightly more of the viral protein NS4b was produced in Huh7.5 cells than in Huh7 cells upon 48 hours of Asibi infection (**Fig. 1G**), which is consistent with the RT-qPCR analysis (**Fig. 1D**). These experiments indicate that both IRF3 and NF-kB are activated during Asibi infection of human hepatoma cells and that RIG-I contributes to IRF3 activation.

Immunofluorescence analysis revealed that IRF3 was detectable exclusively in the cytosol of 100% of uninfected cells, whereas it was found both in the cytosol and the nucleus of around 50% of cells infected for 24 hours (**Fig. 1H**). Infected cells were identified with antibodies recognizing dsRNA structures, including probably viral replicative intermediates (**Fig. 1H**). Similar quantification showed that around 3% of uninfected cells exhibited nuclei positive for the NF-kB subunit p65 whereas around 65% of Asibi-infected cells showed nuclei positive for p65 (**Fig. 1I**).

Together, these results suggest that both the NF-κB and the IRF3 pathways are activated upon Asibi infection of human hepatoma cells and their initiation is dependent on RIG-I.

### RIG-I localizes in stress granules in human and monkey cells infected with YFV

Having determined that RIG-I is important for mediating a cytokine response in Huh7 cells infected with Asibi (**Fig. 1**), we then investigated its subcellular localization. The distribution of the RIG-I signal was found to be weak and diffuse in the cytosol of uninfected Huh7 cells (**Fig. 2A**), which is consistent with data reported for unstimulated HeLa cells (20). After infection with Asibi for 24 hours, RIG-I accumulated in cytosolic foci (**Fig. 2A**). Since RIG-I was previously shown to localize in stress granules in HeLa cells infected with several types of viruses (20–23), the Huh7 cells were also stained with the stress granule marker TIA-1 related protein (TIAR). In non-infected cells, the TIAR signal was weak and diffused in the cytoplasm and in the nucleus of the cells. In infected cells, TIAR was redistributed into cytoplasmic foci (**Fig. 2A**). The RIG-I structures present in infected cells co-localized with TIAR foci (**Fig. 2A**, zoom images for details), suggesting that RIG-I is recruited to stress granules upon infection. The intensity profiles of individual foci confirmed that the RIG-I signal and the TIAR signal overlapped (**Fig. 2A**, right panels). These profiles also revealed that dsRNAs seemed to be excluded from stress granules (**Fig. 2A**).

**Fig. 2:**
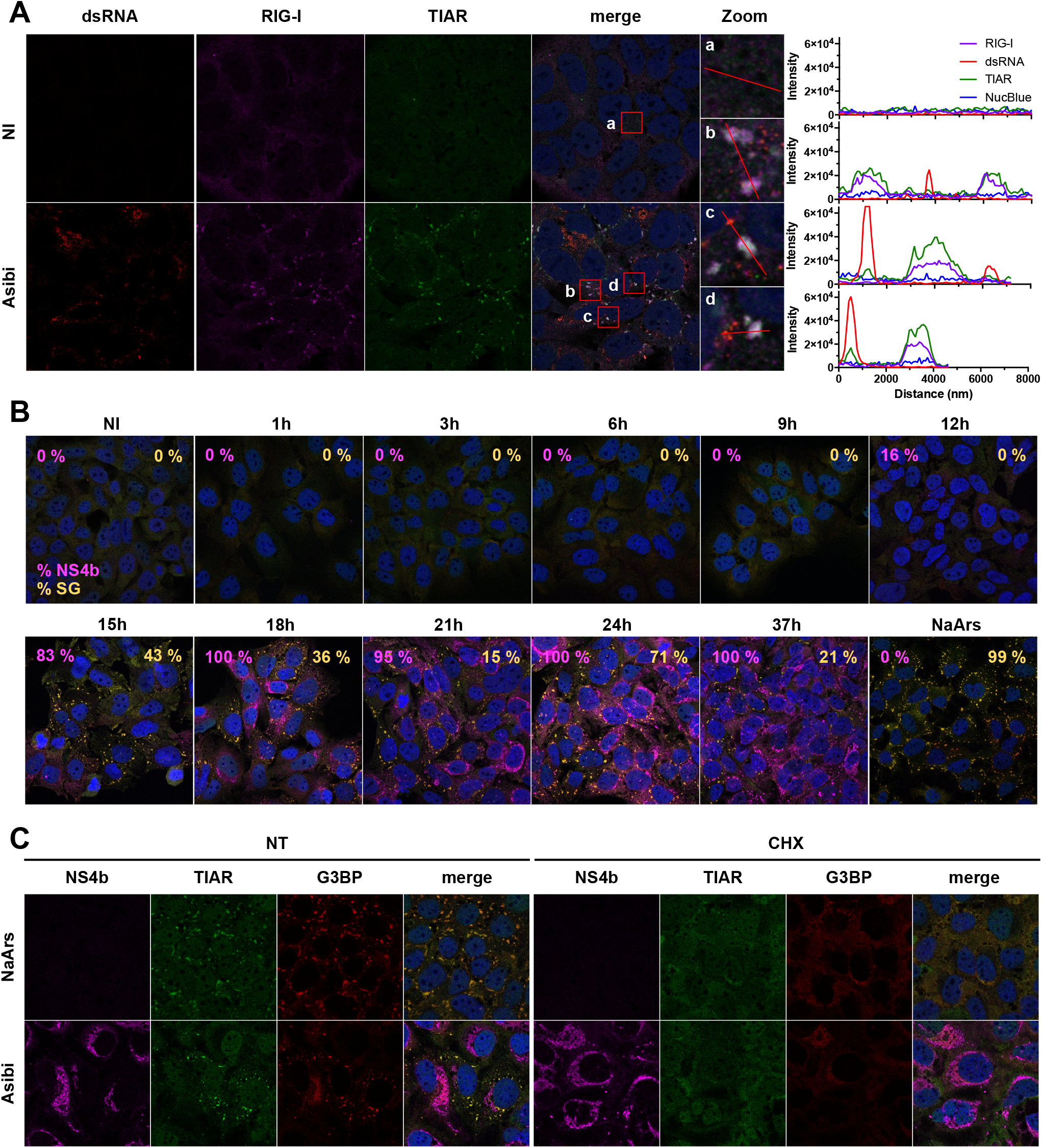
RIG-I localizes to stress granules in Huh7 cells infected with YFV. (**A**) Huh7 cells were left uninfected (NI) or infected with Asibi at MOI of 20 for 24 hours. Cells were then stained with dsRNA (red), RIG-I (purple), TIAR (green) antibodies and NucBlue^®^ (blue). (**B**) Huh7 cells were left uninfected (NI), treated with 0.5 mM NaArs for 30 min or infected with Asibi at an MOI of 20 for the indicated time. Cells were then stained with NS4b (purple), G3BP (red), TIAR (green) antibodies and NucBlue^®^ (blue). Percentage of NS4b-positive and G3BP-positive cells were estimated and shown in purple and yellow respectively. (**C**) Huh7 cells were treated with 0.5 mM NaArs for 30 min or infected with YFV-Asibi at MOI of 20 for 48 hours. Cells were then left untreated (NT) or treated with 100 μg/mL Cycloheximide (CHX) for 1 hour and stained with antibodies against NS4b (purple), TIAR (green) and G3BP (red), as well as with NucBlue^®^ (blue).

To follow the formation of stress granules in infected cells, the localization of TIAR and RAS GTPase-activating protein SH3 domain-binding protein 1 and 2 (G3BP), which are canonical stress granule markers (29), were examined in Huh7 cells infected with Asibi for over time (**Fig. 2B**). Infected cells were identified by the presence of the viral protein NS4b (**Fig. 2B**). TIAR and G3BP were diffuse in non-infected cells. They remained diffuse until 12 hpi, where 16% of cells were determined positive for NS4b. Fifteen hours after infection, around 80% of cells were infected and around 40% of the total number of cells exhibited TIAR- and G3BP-positive foci (**Fig. 2B**). At 24 hpi, all cells were infected and around 70% of them were positive for TIAR- and G3BP-containing foci. At 37 hpi, all cells remained infected and around 20% of them exhibited TIAR- and G3BP positive foci. These foci resembled the ones detected in Huh7 cells treated with sodium arsenite (NaArs), a chemical commonly used for induction of oxidative stress in eukaryotic cells, for 30 minutes (**Fig. 2B**). These results indicate that upon Asibi infection, TIAR- and G3BP-positive foci appeared between 12 and 15 hpi, peak around 24 hpi and persist in around 20% of cells until later times. Moreover, these data suggest that the formation of the stress granules occur throughout infection with Asibi.

To further validate the nature of the TIAR- and G3BP-positive foci observed in infected cells, we assessed their sensitivity to cycloheximide. Cycloheximide is a compound that inhibits stress granule formation and fosters their disassembly by stabilizing mRNAs on polysomes (34). Huh7 cells treated with NaArs for 30 min and subsequently incubated with cycloheximide for 1 hour or left untreated were used as positive controls (**Fig. 2C**). Alternatively, cells were infected with Asibi for 48 hours and then treated, or not, with cycloheximide for 1 hour. The TIAR and the G3BP signals overlapped, both in cells infected by Asibi for 48 hours and in NaArs-treated cells (**Fig. 2C**). Cycloheximide treatment led to a complete loss of TIAR- and G3BP-positive foci, both in NaArs-treated cells and in Asibi-infected cells (**Fig. 2C**, compare non-treated and treated cells). These results suggest that, under both conditions, most mRNAs trapped in these cytoplasmic foci can be released within 1 hour and undergo some translation on polysomes. Most importantly, these experiments confirmed that the cytoplasmic foci containing TIAR, G3BP and RIG-I identified in infected cells also contained mRNAs, and as such, can be defined as canonical stress granules. Furthermore, these experiments also showed that the stress granule proteins TIAR and G3BP are not recruited to Asibi replication complexes identified by NS4b, in contrast to observations reported for ZIKV, DENV and WNV (35, 36).

We next monitored the formation of stress granules upon Asibi infection in different cell models than Huh7. Like Huh7 cells, HepG2 cells are hepatocyte-derived cells and are thus relevant for infection with YFV, a hepatotropic virus. TIAR- and G3BP-positive foci were detected in HepG2 cells infected for 48 hours (**Fig. 3A**). Stress granules were also detected in CMMT cells, a *Macaca mulatta* epithelial cell line, upon 24 hours of infection with Asibi (**Fig. 3B**). Macaques are natural hosts of YFV during the jungle cycle of transmission (37). Similarly, TIAR- and G3BP-positive foci were observed in Huh7 cells infected with the clinical isolate HD1279 (HD Dakar) for 24 hours (**Fig. 3C**). These data indicate that infection with two YFV strains triggers the formation of stress granules in several mammalian cell lines, suggesting that the stress response is neither viral strain nor cell type specific.

**Fig. 3:**
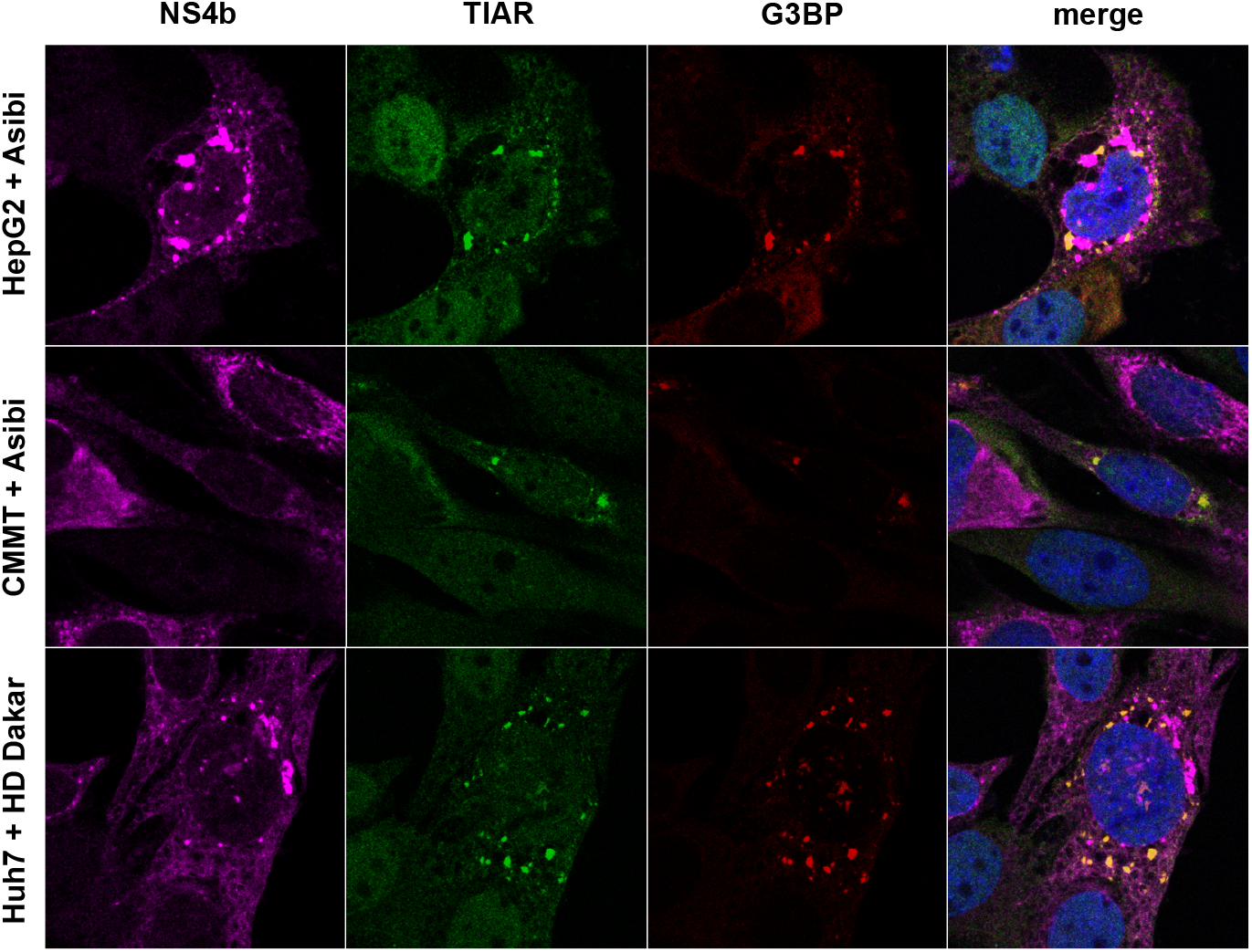
YFV infection induces the formation of stress granules in several mammalian cells. HepG2 or CMMT cells were infected with Asibi at MOI of 20 for, respectively, 48 and 24 hours. Huh7 cells were infected with YFV-HD Dakar at MOI of 10 for 24 hours. Cells were then stained with NS4b (purple), TIAR (green), G3BP (red) antibodies and NucBlue^®^ (blue).

### PKR contributes to stress granule formation and innate response in Asibi-infected Huh7 cells

PKR is known to be activated upon infection with several flaviviruses, including ZIKV, WNV and Japanese Encephalitis virus (JEV) (38–40) and to play an important role in stress granule nucleation through its ability to phosphorylate the eukaryotic initiation factor eIF2α (41). We first assessed whether PKR was activated upon Asibi infection of Huh7 cells. Western blot analysis showed that PKR was phosphorylated in Huh7 cells infected for 24 and 48 hours, but not in non-infected cells (**Fig. 4A**). Thus, PKR is activated upon Asibi infection of Huh7 cells. To evaluate the role of PKR in the formation of stress granules in Asibi-infected human hepatic cells, we generated a Huh7 short hairpin (sh)-PKR cell line by transduction with lentiviruses expressing short hairpin RNAs (shRNAs) specific for PKR. While PKR expression was detectable in Huh7 cells transduced with lentivirus expressing control shRNA targeting luciferase (Huh7-shLuc cells), it was not detectable in Huh7-shPKR cells (**Fig. 4B**). As a control experiment, we treated the cells with NaArs, which induces stress granule formation *via* HRI-mediated eIF2α phosphorylation (42). TIAR- and G3BP-positive stress granules were detected in both Huh7-shLuc and Huh7-shPKR cells treated with NaArs for 30 minutes (**Fig. 4C**). This confirms that the absence of PKR does not disrupt the formation of NaArs-dependent stress granule induction.

Automated microscopic quantification revealed that around 10% of Huh7-shLuc cells infected at a MOI of 2 for 24 or 34 hours contained at least two stress granules (**Fig. 4 D-E**). At a MOI of 20, around 25% of Huh7-shLuc cells were positive for stress granules at 24 and 34 hpi (**Fig. 4 D-E)**. No TIAR-nor G3BP-positive foci were detected in Huh7-shPKR cells infected at a MOI of 2 for 24 or 34 hours. At a MOI of 20, a low percentage (around 1%) of Huh7-shPKR cells exhibited detectable stress granules (**Fig. 4 D-E)**. These stress granules possibly formed in a PKR-independent manner, *via* the activation of other members of the eIF2α kinase family. Alternatively, a minute quantity of PKR still being expressed in shPKR cells may initiate the formation of these granules. A similar proportion of Huh7-shLuc and Huh7-shPKR cells were found positive for NS4b at 24 or 34 hpi at the two tested MOIs (**Fig. 4F**). Together, we conclude that PKR plays a key role in stress granule formation in Asibi-infected human hepatic cells and that the absence of stress granules has no effect on viral infection.

**Fig. 4:**
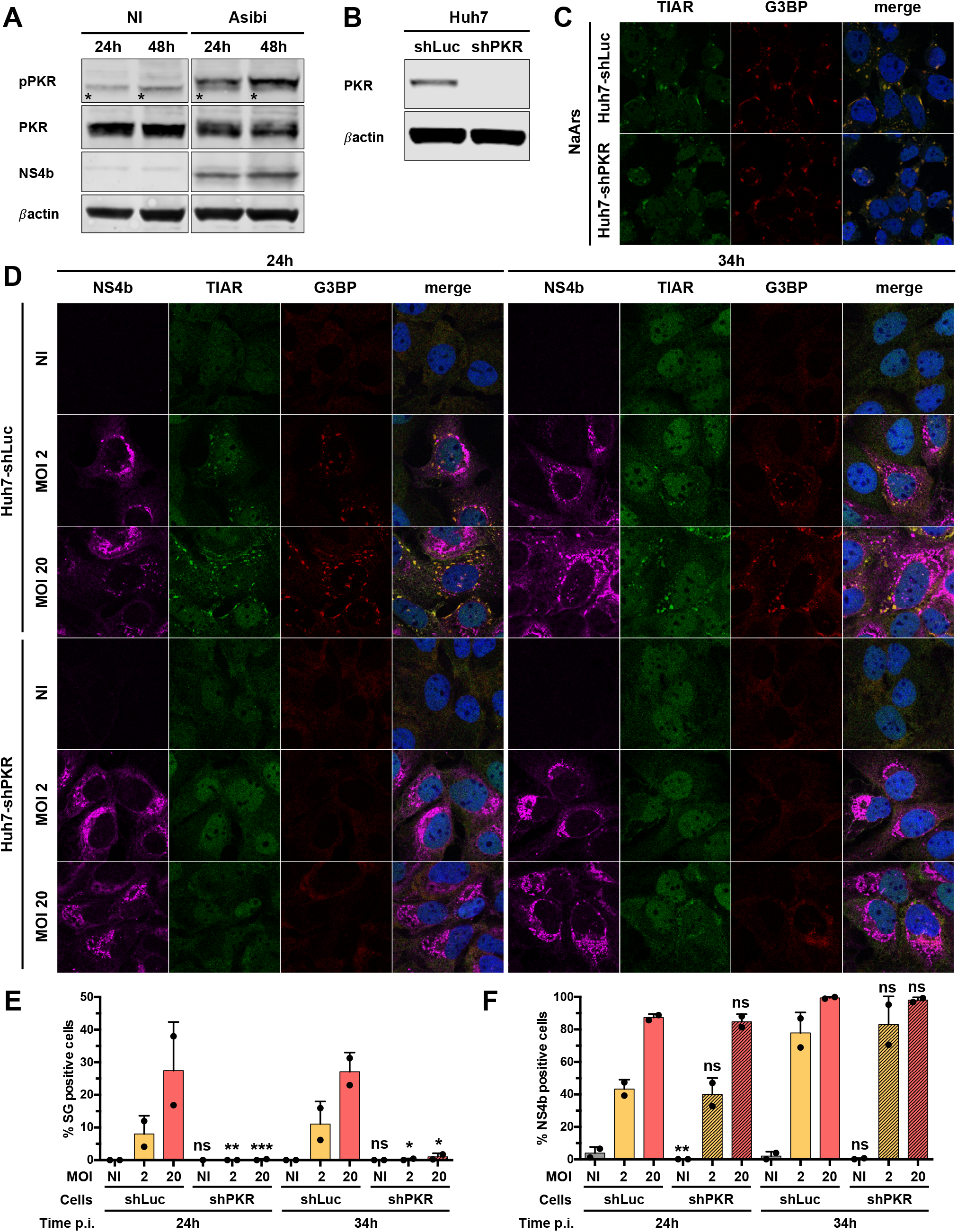
PKR contributes to stress granule formation in Asibi-infected Huh7 cells. (**A**) Huh7 cells were left uninfected (NI) or infected with Asibi at an MOI of 20 for 24 or 48 hours. Whole-cell lysates were analysed by Western blotting with antibodies against the indicated proteins. Samples were loaded on the same gel. *: unspecific bands. (**B**) Whole-cell lysates of Huh7-shLuc and Huh7-shPKR cells were analysed by Western blotting with antibodies against the indicated proteins. (**C**) Huh7-shLuc and Huh7-shPKR cells were treated with 0.5 mM NaArs for 30 min and stained with TIAR (green) and G3BP (red) antibodies, as well as with NucBlue^®^ (blue). (**D**) Huh7 cells were left uninfected (NI) or infected with YFV-Asibi at MOI of 2 or 20 for 24 or 34 hours. Cells were then stained with NS4b (purple), TIAR (green), G3BP (red) antibodies and NucBlue^®^ (blue). Images are representative of two independent experiments. Percentage of G3BP-positive (**E**) or NS4b-positive cells (**F**) were estimated by analysing at least 200 cells per condition. ns: non-significant, *: p<0.05, **: p<0.01, ***: p<0.001.

Since no stress granules were detected in cells depleted of PKR in three of four conditions tested (**Fig. 4E**), we used Huh7-shPKR cells to determine whether absence of granules affected cytokine induction. Consistent with our Western blot data (**Fig. 4B**), RT-qPCR analysis showed that *PKR* mRNAs abundance was reduced by more than 90% in Huh7-shPKR cells in comparison to Huh7-shLuc cells (**Fig. 5A**). (**Fig. 5B**), which is consistent with our microscopic analysis (**Fig. 4F**). *IL6* mRNA abundance was significantly lower in Huh7-shPKR cells than that of Huh7-shLuc when cells were infected at a MOI of 20 (**Fig. 5C**). The abundances of *TNFA* and *IFNB* mRNAs were significantly lower in Huh7-shPKR cells than those in infected Huh7-shLuc under the same conditions of infection (**Fig. 5D-E**). This is consistent with our preceding observation of cytokine induction in Asibi infected cells being dependent on NF-kB (**Fig. 1**) and with a previous report showing that PKR is involved in the induction of the *IFN* through the activation of NF-kB (44). Thus, at least two nucleic acid sensors, namely RIG-I (**Fig. 1**) and PKR (**Fig. 4** and **5**), contribute to the antiviral response against Asibi in Huh7 cells.

**Fig. 5:**
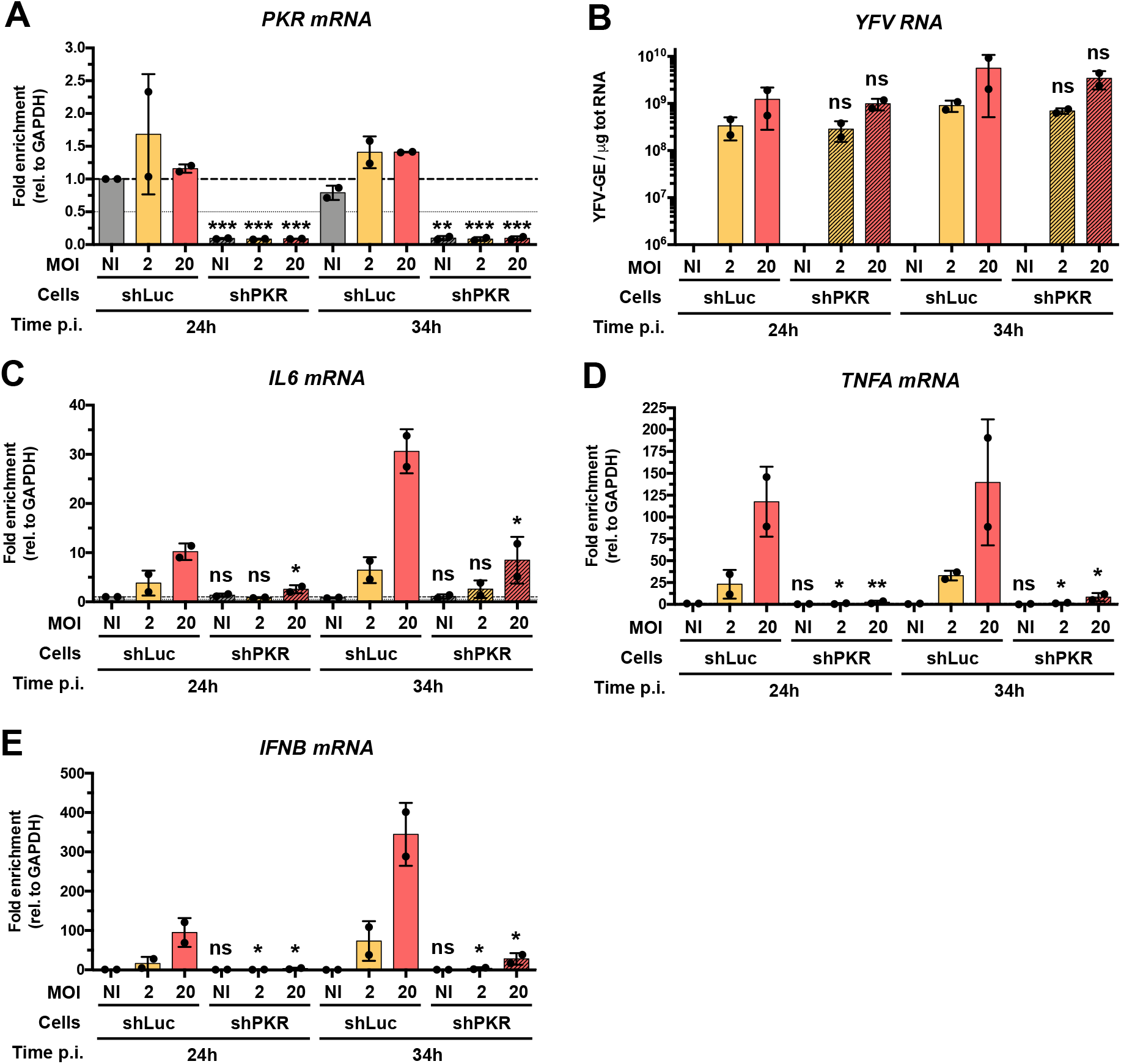
PKR contributes to innate response in Asibi-infected Huh7 cells. Huh7-shLuc or -shPKR cells were left uninfected (NI) or infected with Asibi at MOI of 2 or 20 for 24 or 34 hours. The relative amounts of (**A**) *PKR* mRNA, (**C**) *IL6* mRNA, (**D**) *TNFA* mRNA and (**E**) *IFNB* mRNA were determined by qPCR analysis and normalized to that of GAPDH mRNA and non-infected shLuc samples. The relative amounts of cell-associated viral RNA were determined by RT-qPCR analysis (**B**) and expressed as genome equivalents (YFV-GE) per μg of total cellular RNA. Mean ± SD is represented. One-way ANOVA on log transformed data with Tukey corrected multiple comparisons tests were performed. shPKR samples were compared to shLuc ones. ns: non-significant, *: p<0.05, **: p<0.01, ***: p<0.001.

Taken together, our data show that PKR contributes to both stress granule formation and induction of cytokines in Huh7 cells infected with Asibi. However, since PKR does not require its kinase activity to stimulate NF-κB (44, 45), it might contribute to the cytokine induction in Asibi-infected cells in a eIF2α- (and thus stress granule-) independent manner.

### Induction of pro-inflammatory cytokines is not affected by the inhibition of stress granules

To further investigate the putative link between stress granules and the induction of antiviral cytokines, we used a siRNA-knockdown approach to inhibit stress granule assembly. We tested numerous combinations of siRNAs targeting core granule components in infected cells. The most efficient strategy consisted in transfecting Huh7 cells twice with four pools of siRNAs targeting TIAR, TIA-1, G3BP1 and G3BP2 (siSG). In these cells, we monitored stress granule formation using antibodies against eIF3b and eIF4G, two core components of stress granules. Around 30% of Huh7 cells transfected with scrambled control siRNAs and infected at a MOI of 2 or 20 for 24 hours contained at least two eIF4G- and eIF3b-positive foci (**Fig. 6A-B**). When cells were transfected with the 4 siRNA pools and infected under the same conditions and for the same amount of time, less than 5% of cells exhibited stress granules (**Fig. 6A-B**). The impact of the siRNA against TIAR, TIA-1, G3BP1 and G3BP2 on the number of cells positive for stress granules following 34 hours of infection was also significant at the two tested MOIs (**Fig. 6B**). Under the four tested conditions, the surface area occupied by stress granules per cell was significantly reduced in cells transfected with the mixture of four siRNA pools compared to the controls (**Fig. 6C**). These experiments validate our silencing approach as an efficient way to significantly reduce both the number of cells positive for stress granules and the surface occupied by stress granules per cell. Transfection with the 4 siRNA pools had no effect on viral infection, when compared to transfection with control siRNAs (**Fig. 6D**). RT-qPCR analysis showed that TIAR, TIA-1, G3BP1 and G3BP2 mRNA abundances were significantly reduced in cells transfected with the 4 siRNA pools as compared to cells transfected with control siRNAs (**Fig. 7A-D**). Of note, TIAR and TIA-1 expression was induced three- to four-fold by infection, in an MOI-dependent response. This may be a consequence of virally-induced stress. Transfection of the four siRNA pools has no effect on *TNFA, IL6* and *IFNB* mRNA abundances nor on viral RNA yield (**Fig. 7E-H**). Together, these data show that significant reduction of stress granule numbers and size did not suppress the cytokine response to Asibi infection. Thus, in our model, stress granules do not contribute to the initiation of the antiviral response.

**Fig. 6:**
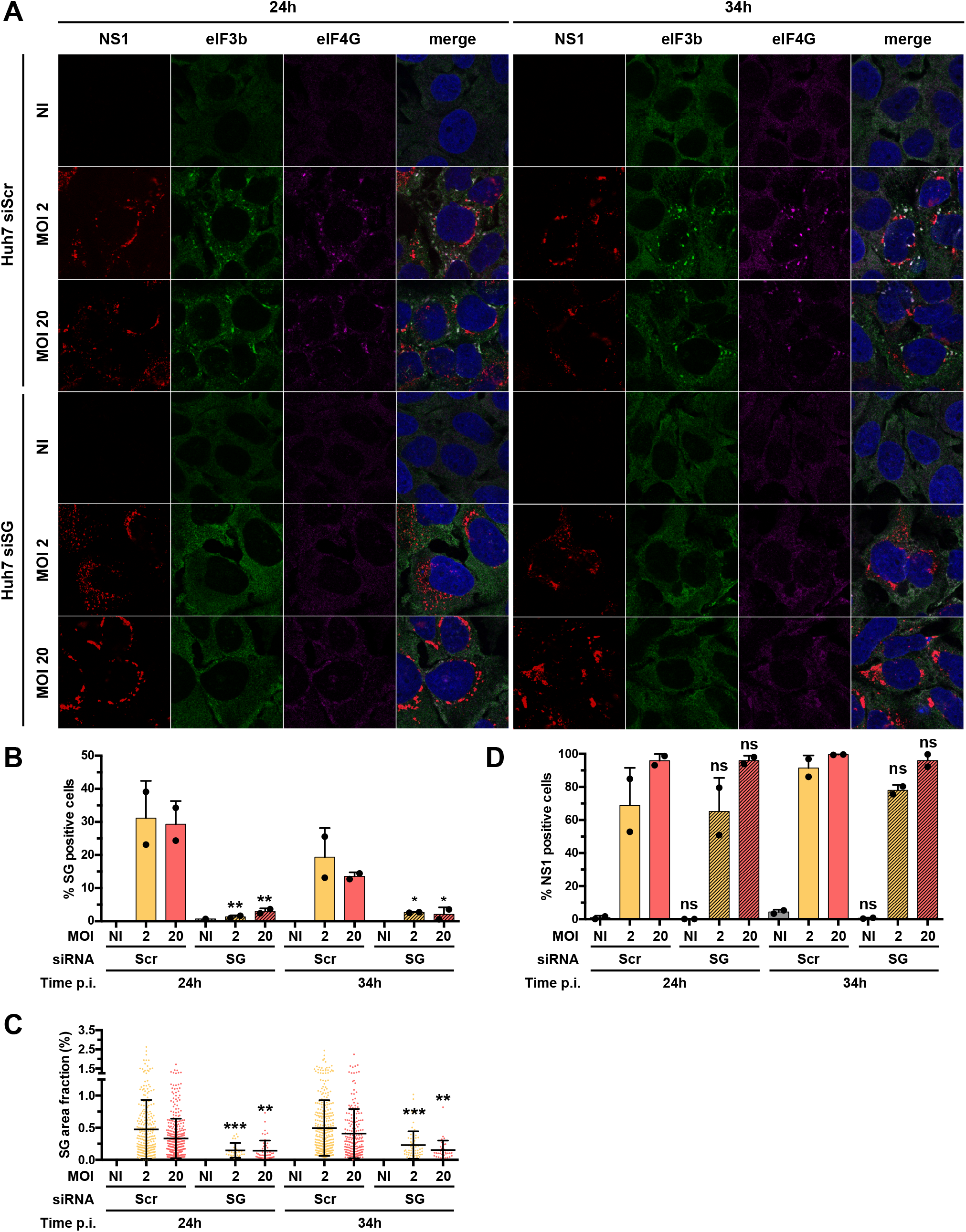
Knock-down of G3BP1, G3BP2, TIAR and TIA1 inhibits stress granules formation. (**A**) Huh7 cells were transfected with scramble siRNA (siScr) or with siRNA targeting G3BP1, G3BP2, TIAR and TIA1 (siSG) 48 and 24 hours prior to Asibi infection. Transfected cells were left uninfected (NI), treated with 0.5 mM NaArs for 30 min (NaArs) or infected with Asibi at MOI of 2 or 20 for 24 or 34 hours. Cells were then stained with NS1 (purple), eIF3b (green), eIF4G (red) antibodies and NucBlue^®^ (blue). Images are representative of two independent confocal microscopy analyses. Percentages of eIF4G- and eIF3b-positive (**B**) or NS1-positive cells (**D**) were estimated. (**C**) Stress granule area was determined for each cell as the sum of the whole area occupied by stress granules divided by the total cell surface. (**B-D**) Around 200 cells per independent experiment were scored. ns: non-significant, *: p<0.05, **: p<0.01, ***: p<0.001.

**Fig. 7:**
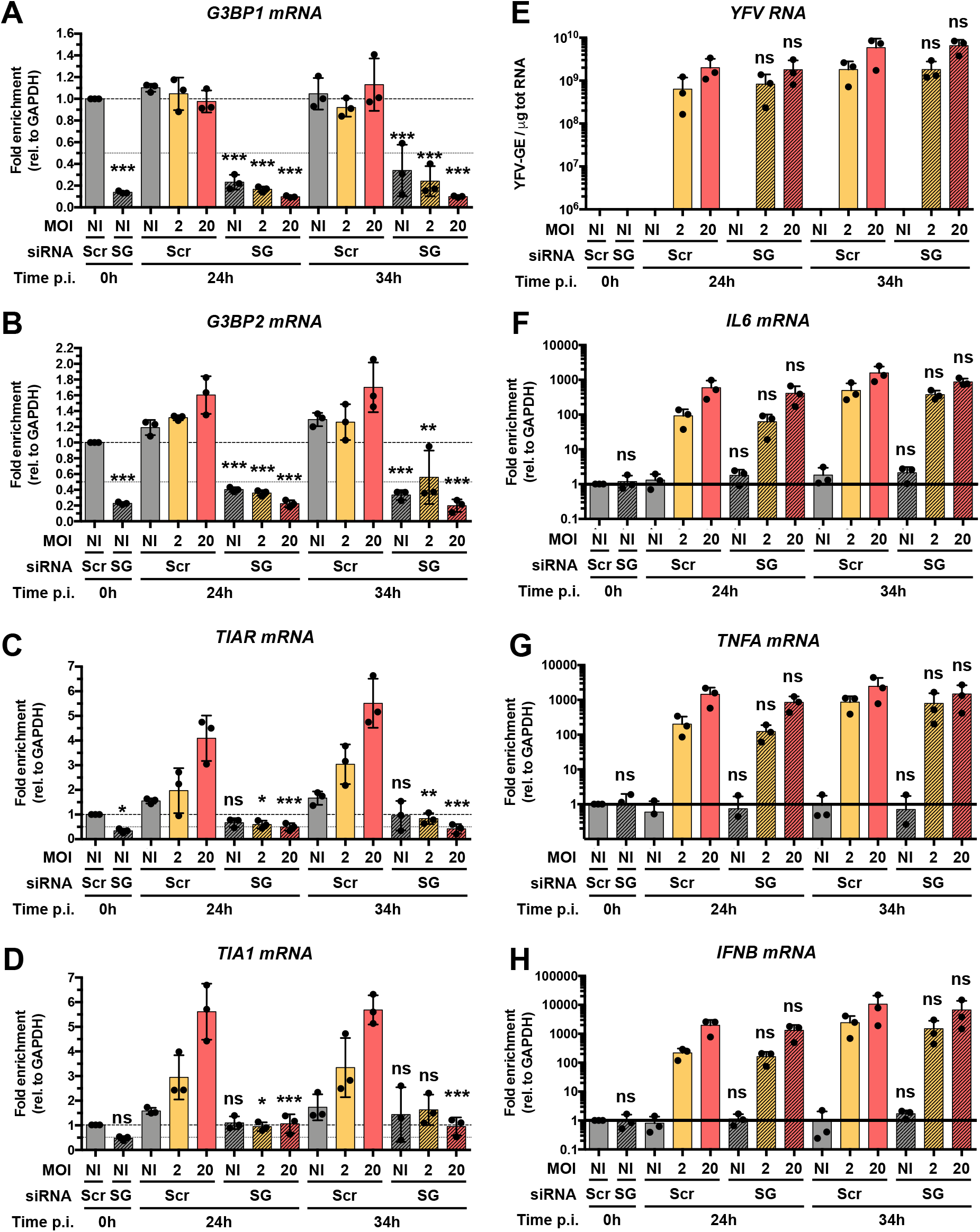
Inhibition of stress granule formation has limited effect on antiviral response. Huh7 cells were transfected with scramble siRNA (siScr) or with four pools of siRNA targeting G3BP1, G3BP2, TIAR and TIA1 (siSG) 48 and 24 hours prior to infection. Transfected cells were left uninfected (NI) or infected with Asibi at an MOI of 2 or 20. Cells were harvested prior to infection (0 h), 24 or 34 hours post-infection. The relative amount of (**A**) *G3BP1* mRNA, (**B**) *G3BP2* mRNA, (**C**) *TIAR* mRNA, (**D**) *TIA1* mRNA, (**F**) *IL6* mRNA, (**G**) *TNFA* mRNA and (**H**) *IFNB* mRNA was determined by qPCR analysis and normalized to that of GAPDH mRNA and non-infected sh-Scr samples. The relative amounts of cell-associated viral RNA were determined by RT-qPCR analysis (**E**) and expressed as genome equivalents (YFV-GE) per μg of total cellular RNA. Mean ± SD is represented. One-way ANOVA on log transformed data with Tukey corrected multiple comparisons test were performed. Samples treated with siSG-RNA were compared to siScr samples. ns: non-significant, *: p<0.05, **: p<0.01, ***: p<0.001.

### RIG-I signaling is not involved in stress granule assembly

In Huh7 cells, RIG-I signaling is crucial for cytokine induction and secretion upon Asibi infection (**Fig. 1**). Therefore, we wondered whether RIG-I signaling would participate in stress granule assembly and/or maintenance. We quantified the proportion of Huh7 and Huh7.5 cells that were positive for stress granules at 24 and 34 hpi following infection at two MOIs. A similar proportion of Huh7 and Huh7.5 cells were found positive for stress granule markers in all tested conditions (**Fig. 8 A-B**). No difference in the number of cells positive for NS4b was observed between Huh7 and Huh7.5 cells (**Fig. 8C**). We conclude that the absence of RIG-I signaling had no effect on the number of granules that assemble in response to infection. It is thus unlikely that RIG-I mediates a feed-forward loop to maintain stress granule assembly.

**Fig. 8:**
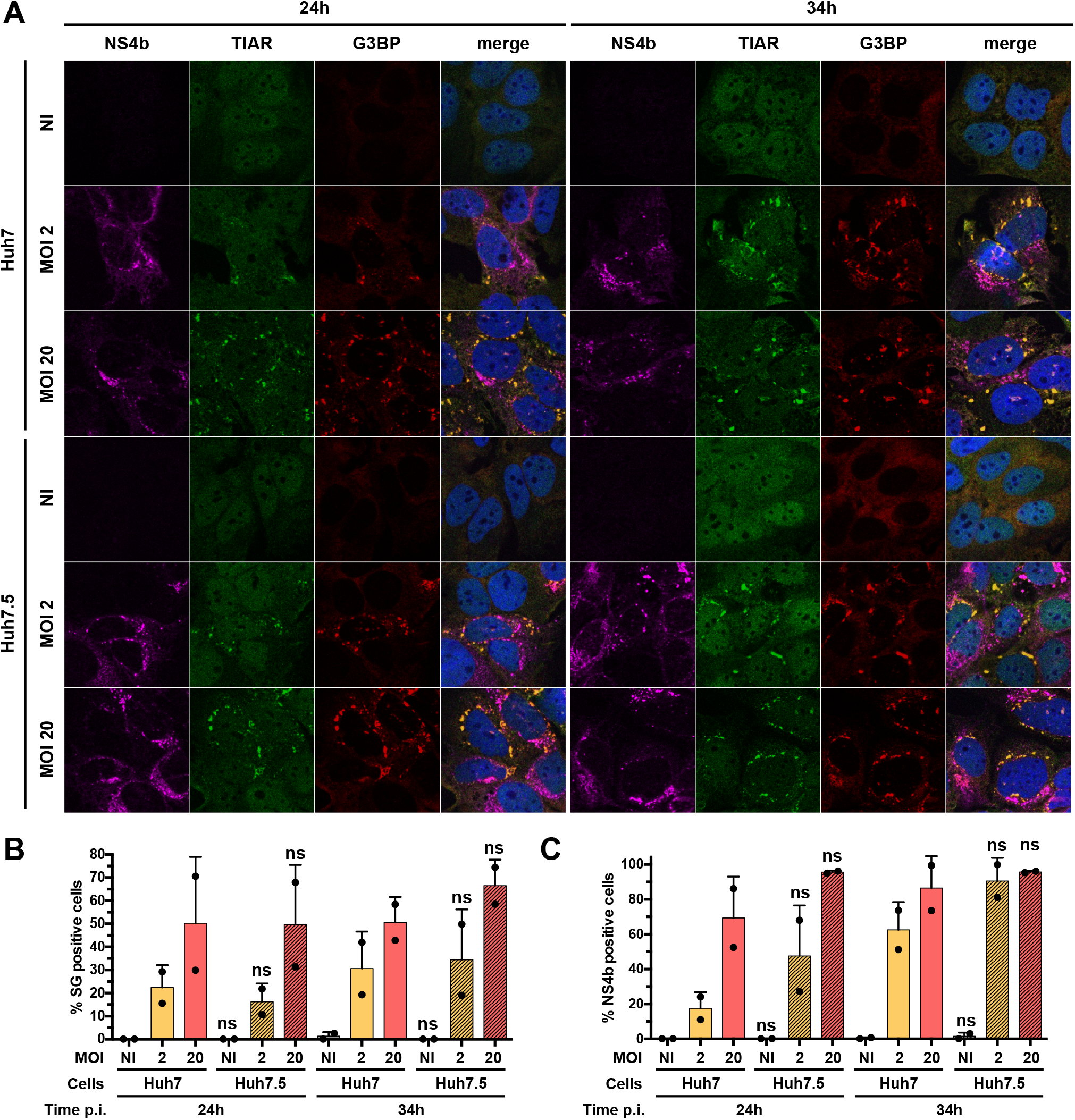
RIG-I signaling is not involved in stress granule assembly. (**A**) Huh7 and Huh7.5 cells were left uninfected (NI), treated with 0.5 mM NaArs for 30 min (NaArs) or infected with Asibi at MOI of 2 or 20 for 24 or 34 hours. Cells were then stained with NS4b (purple), TIAR (green), G3BP (red) antibodies and NucBlue^®^ (blue). (**B-C**) Images are representative of two independent confocal microscopy analyses. Around 200 cells per independent experiment were scored. Percentages of G3BP-positive (**B**) or NS4b-positive cells (**C**) were estimated. ns: non-significant, *: p<0.05, **: p<0.01, ***: p<0.001.

### YFV RNA is excluded from stress granules

Our previous analysis showed that neither dsRNA (**Fig. 2A**) nor the viral proteins NS4b and NS1 (**Fig. 2B, 2C, 3, 4D, 6A** and **8A**) seem to be recruited to stress granules. To further investigate the putative association between stress granules containing RIG-I (**Fig. 2A**) and viral RNA, we used an RNA fluorescence *in situ* hybridization (FISH) approach coupled with immunofluorescence and confocal microscopy. The Asibi-specific FISH probe produced no signal in non-infected Huh7 control cells (**Fig. 9A**) and a bright punctate signal in the cytoplasm of infected cells (**Fig. 9A**), confirming the specificity of the probe (46). Asibi RNA was never found to be associated with TIAR (**Fig. 9A**). This was unexpected since RIG-I signaling is key to cytokine induction and production (**Fig. 1**) and massively concentrate in stress granules in Asibi-infected cells (**Fig. 2A**). These results suggest that an interaction between viral RNA and RIG-I might not happen in stress granules in Asibi-infected Huh7 cells. A minute quantity of RIG-I might however interact with viral RNA in ER-derived replication sites and initiates a stress granule-independent IFN response. In line with this, RIG-I was recruited to TIAR-positive granules in Huh7 cells treated with NaArs (**Fig. 9B**), which is consistent with previous observations on NaArs-treated HeLa cells (20). These data further indicate that RIG-I recruitment to stress granules can occur in absence of viral RNA.

**Fig. 9:**
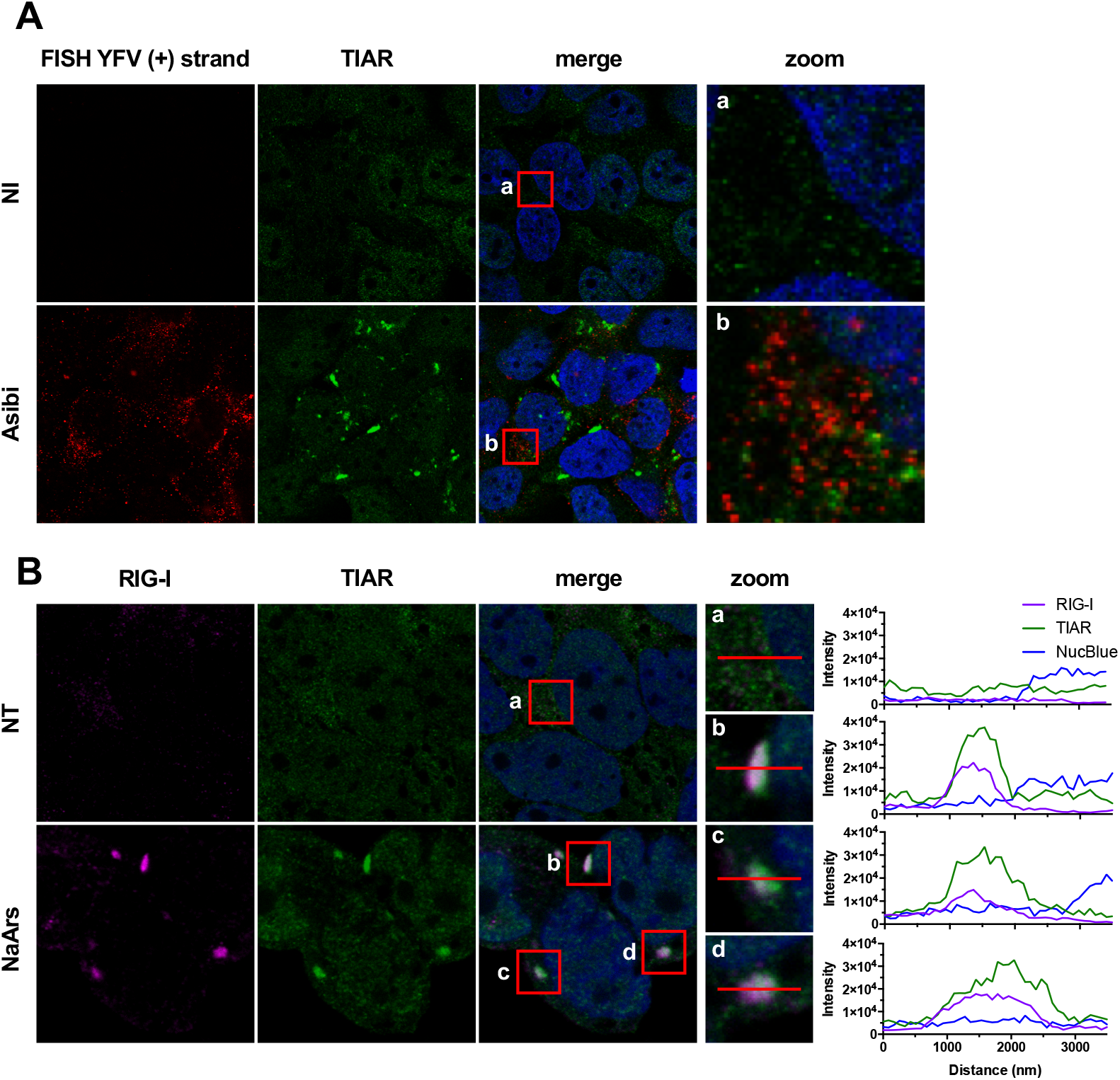
YFN RNA is excluded form stress granules. (**A**) Huh7 cells were left uninfected (NI) or infected with Asibi at MOI of 20 for 24 hours. Cells were then stained with TIAR (green) and NucBlue^®^ (blue), prior to be processed for FISH using a probe specific for YFV (+) strand RNA (red). (**B**) Huh7 cells were left untreated (NT) or treated with 0.5 mM NaArs for 30 min. Cells were then stained with RIG-I (purple), TIAR (green) antibodies and NucBlue^®^ (blue).

## Discussion

The identity of the PRR(s) responsible for the initiation of a pro-inflammatory response upon direct infection of human hepatocytes with wild-type strains of YFV has not been uncovered thus far. In this study, we first identified RIG-I and PKR as main actors responsible for the induction and production of pro-inflammatory cytokines upon infection of human hepatocytes with the Asibi strain of YFV. We demonstrated the crucial involvement of RIG-I in this process by directly comparing the dysfunctional RIG-I-expressing human hepatoma cell line Huh7.5. with its RIG-I-competent counterpart (Huh7 cells). The level of IL6 and TNFα secreted in the first 48 hours of infection by Huh7.5 cells upon Asibi infection was under the limit of detection, whilst both cytokines were produced in Huh7 cells in the same experimental conditions. This is in accordance with our previous studies showing that human plasmacytoid dendritic cells infected with the YFV vaccine strain 17D induced an IFN response in a RIG-I dependent manner (19). Other members of the RLR family may contribute to YFV sensing in other cell types or at later time of infection. Such distinct temporal activity of RIG-I and MDA5 during infection with the related West Nile Virus has been reported in mice (47).

In Huh7 cells silenced for PKR and infected with Asibi, cytokine expression was, in most of the tested conditions, below the detection limit. This suggests that PKR, like RIG-I, largely contributes to the initiation of the antiviral response. This is consistent with the previously reported role of PKR in the production of IFN and other pro-inflammatory cytokines upon viral infection, possibly *via* its interaction with the IKK complex and subsequent activation of NF-κB (44, 48). Moreover, PKR was previously described as an important effector protein against infection with WNV in mice (40).

The fact that abolishing RIG-I signaling or silencing PKR drastically reduces the pro-inflammatory response in Asibi infected cells shows the non-redundant nature of these two dsRNA sensors in maintaining cytokine production. We speculate that RIG-I and PKR cooperate to induce an anti-YFV response. Indeed, PKR interacts with RIG-I and MDA5 in HEK293T cells transfected with FLAG-tagged versions of RIG-I or MDA5 and infected with a mutant vaccinia virus lacking a gene known to inhibit PKR (49). However, in this context, PKR was only required for MDA5, but not RIG-I mediated, IFN production (49). Consistent with the hypothesis that PKR and RIG-I cooperate to orchestrate antiviral response, PKR is known to interact with several members of the TRAF family, including TRAF2 and TRAF6 (50, 51), two proteins involved in MAVS signaling. Finally, MAVS, which acts downstream of RIG-I and MDA5, interacts with PKR (52) and is essential for PKR-induced IFN production (49).

Stress granules have been demonstrated to contain and concentrate viral nucleic sensors; hence, they are generally regarded as a functional platform for ligand recognition and subsequent induction of an antiviral response. We report here that Asibi replication triggers the formation of stress granules in human hepatoma cells and epithelial monkey cells. This is consistent with previous studies showing that infection with the vaccine strain 17D induces the formation of stress granules in human dendritic cells and hamster kidney cells (53). Thus, formation of stress granules seems to be a general feature of flavivirus infection (35, 36, 54, 55). However, unlike WNV, DENV, ZIKV and the related tick-borne encephalitis virus (35, 36, 38, 54–57), YFV does not seem to efficiently antagonize stress granule biogenesis. YFV may not express proteins that inhibit granule formation. Alternatively, stress granules may not have an antiviral function against YFV and thus the virus has no pressure to evolve an evasion strategy.

Stress granules were not detected in Huh7 cells depleted of PKR upon 24 hours of Asibi infection, strongly suggesting a PKR-dependent granule establishment early in infection. Of note, RIG-I signaling was not involved in stress granule assembly, indicating that, whilst the two sensors are both critically implicated in the pro-inflammatory cytokine production, only PKR is able to activate stress granule formation. At 34 hours, very few cells silenced for PKR exhibited stress granules. Small quantities of remaining PKR, below the detection limit of Western blot and qPCR analysis, may be responsible for the formation of these foci. Alternatively, they may form subsequent to the phosphorylation of eIF2α by one of the other three eIF2α kinases. YFV replication and assembly, which takes place within ER membranes (3), likely causes an ER stress and might thus trigger the activation of the ER resident kinase PERK. Since GCN2 is phosphorylated in mouse and human dendritic cells upon infection with 17D (53), this kinase could also potentially contribute to granule formation in Asibi infected cells.

Similar to what was reported in the context of infection with numerous RNA viruses (27, 28, 58), we showed that RIG-I accumulates in stress granules upon Asibi infection. PKR, MDA5, LGP2 and the DNA sensor cGAS were also shown to concentrate in stress granules in cells infected with a large panel of RNA and DNA viruses (20, 22–26). However, we were unable to detect neither Asibi RNA nor dsRNA structures in stress granules of infected cells. Thus, the interaction between YFV RNA by RIG-I and PKR is unlikely to occur in stress granules. Similarly, in cells infected with Orthohantaviruses, no viral RNA was found associated with stress granules (59). Conversely, viral RNA was detected in stress granules induced by infection with IAVΔNS1 (20). We propose that, at least in Asibi-infected cells, the large accumulation of PKR and RIG-I in stress granules masks the detection of small quantities of PKR and RIG-I which are recruited to perinuclear ER-derived replication sites, where viral RNA sensing may occur. Importantly, inhibition of stress granule formation didn’t affect the pro-inflammatory cytokine expression by infected hepatocytes, indicating that the anti-viral response and stress granules formation are independent events in the early phases of YFV infection. Whether the formation of stress granules and their composition affect later stages of infection remains to be established.

Collectively, our data strongly suggest that, in cells infected with YFV, pro-inflammatory cytokine induction is mediated by RIG-I and PKR, independent of their localization in stress granules. Thus, contrary to the widely believed consensus, we demonstrated that the stress granule formation can be uncoupled from the initiation of a pro-inflammatory response upon viral infection.

## Materials and Methods

### Cell lines, virus, antibodies and reagents

Huh7 human hepatocellular carcinoma cells (kindly provided by A. Martin, Institut Pasteur Paris) and Huh7.5 cells (60) were maintained in Dulbecco’s modified eagle’s medium (DMEM)(Gibco) containing GlutaMAX I and sodium pyruvate (Invitrogen) supplemented with 10% heat-inactivated foetal bovine serum (FBS, Dominique Dutscher) and 1% penicillin and streptomycin (P/S, 10.000 U/ml, Thermofischer). HepG2 cells (kindly provided by C. Neuveut, Institut Pasteur Paris) were maintained in DMEM-F12 complemented with 10% FBS, 3.5× 10^-7^ M hydrocortisone, and 5 μg/ml insulin. Human embryonic kidney (HEK) 293FT cells (American Type Culture Collection; ATCC CRL-1573), *Macaca mulatta* mammary gland epithelial CMMT cells (kindly provided by O. Schwartz, Institut Pasteur Paris) and African green monkey kidney epithelial Vero cells (ATCC) were maintained in DMEM 10% FBS.

The Asibi strain, which is the YFV reference strain was provided by the Biological Resource Center of Institut Pasteur. The YFV-DAK strain (YFV-Dakar HD1279) was provided by the World Reference Center for Emerging Viruses and Arboviruses (WRCEVA), through the University of Texas Medical Branch at Galveston, USA. Virus stocks were propagated on Vero cells. Viruses were concentrated by polyethylene glycol 6000 precipitation and purified by centrifugation in a discontinued gradient of sucrose. Infectious virus titers were determined by plaque assays on Vero cells. Cells were infected with 10-fold serial dilutions of viral stocks and incubated for 4 days in DMEM containing 2% FBS and carboxymethyl cellulose. Cells were then washed with PBS, fixed with 3% formaldehyde crystal violet solution for 20 min at room temperature, rinsed with water and plaques were counted. Cell infections were carried out at the indicated MOIs. The viral inoculum was replaced with fresh culture medium two hours post-infection.

The following antibodies were used: pIRF3 S386 (clone EPR2346, Abcam), IRF3 (FL-425, Santa Cruz), NF-κB p65 (D14E12, Cell Signalling), IκBα (L35A5, Cell Signalling), β-Actin (clone AC-74, Sigma-Aldrich), dsRNA J2 (English & Scientific Consulting Kft), G3BP (BD Biosciences), TIAR (C-18, Santa-Cruz), eIF4G (sc-11373, Santa-Cruz), eIF3b (N-20, sc-16377), YFV-NS4b (kind gift from C.M. Rice, Rockefeller University), Alexa Fluor^®^ 488 donkey Anti-Goat IgG (H+L), Alexa Fluor^®^ 594 donkey Anti-Mouse IgG (H+L), Alexa Fluor^®^ 647 donkey Anti-Rabbit IgG (H+L) (Life Technologies), anti-mouse 680 (LI-COR Bioscience), anti-rabbit 800 (Thermo Fisher Scientific). The RIG-I antibody was previously described (20).

Sodium arsenite (NaArs, Sigma) and Cycloheximide (CHX, Sigma) were diluted in water and used at the indicated concentration.

### Generation of shRNA-expressing Huh7 cells by lentiviral transduction

pALPS-shPKR1 and pALPS-shPKR3 were generated by cloning the following sequences in pALPS (61): (TGC TGT TGA CAG TGA GCG CCA GCA GAT ACA TCA GAG ATA ATA GTG AAG CCA CAG ATG TAT TAT CTC TGA TGT ATC TGC TGA TGC CTA CTG CCT CGG A) and (TGC TGT TGA CAG TGA GCG ACA GAA GAA AAG ACT AAC TGT ATA GTG AAG CCA CAG ATG TAT ACA GTT AGT CTT TTC TTC TGG TGC CTA CTG CCT CGG A). HEK293FT cells were transfected with pCMV-VSV-G codon-optimized, pCMVR8.74 (both plasmids were given by P. Charneau, Institut Pasteur) and lentiviral vectors (pALPS-shPKR1/3 or pALPS-shLuc) at a mass ratio of 1:4:4. Huh7 cells were transduced with supernatant from transfected HEK293FT and selected with 10 μg/μL puromycin (Sigma Aldrich) four days later.

### Small Interfering RNA (siRNA) transfection

Huh7 cells were transfected using Lipofectamine RNAiMax (Life Technologies) at 30 nM final of nontargeting scrambled negative control (siScr) or G3BP1, G3BP2 and TIAR siRNAs. Alternatively, cells were transfected at 40 nM final of siRNA-TIA1. All siRNAs were obtained from Dharmacon (ON-TARGETplus SMARTpool). Transfected cells were harvested or used for infection assays 48 hours after transfection.

### Enzyme-linked immunosorbent assay (ELISA)

The amount of TNFα and IL6 in cell-culture supernatants was quantified by ELISA using the Human TNFα or IL6 ELISA kit (eBiosciences), according to manufacturer’s instructions. The detection limits were 4 pg/ml for TNFα and 2 pg/ml for IL6.

### Western blot analysis

Cells were lysed in radioimmunoprecipitation assay (RIPA) buffer (Sigma) containing a protease and phosphatase inhibitor mixture (Roche). Cell lysates were normalized for protein content, boiled in NuPAGE LDS sample buffer (Thermo Fisher Scientific) in non-reducing conditions and the proteins were separated by SDS-PAGE (NuPAGE 4-12% Bis-Tris Gel, Life Technologies). Proteins were transferred to nitrocellulose membranes (Bio-Rad) using a Trans-Blot Turbo Transfer system (Bio-Rad). After blocking with PBS-Tween-20 0,1% (PBST) containing 5% milk or BSA for 1 hour at RT, the membrane was incubated overnight at 4 °C with primary antibodies diluted in a blocking buffer. Finally, the membrane was incubated for 45 min at RT with secondary antibodies (Anti-rabbit/mouse IgG (H+L) DyLight 680/800) diluted in a blocking buffer, washed, and scanned using an Odyssey CLx infrared imaging system (LI-COR Bioscience).

### RNA extraction and RT-qPCR analysis

Total RNAs were extracted from cell-lysates using the NucleoSpin RNA II Kit (Macherey-Nagel) following the manufacturer’s protocol and were eluted in nuclease-free water. First-strand complementary DNA synthesis was performed with the RevertAid H Minus M-MuLV Reverse Transcriptase (Thermo Fisher Scientific). Quantitative real-time PCR was performed on a real-time PCR system (QuantStudio 6 Flex, Applied Biosystems) with SYBR Green PCR Master Mix (Life Technologies). Data were analyzed with the ΔΔCT method, with all samples normalized to GAPDH. All experiments were performed in technical triplicate. Primers used in this study are: GAPDH (GGT CGG AG TCA ACG GAT TTG and ACT CCA CGA CGT ACT CAG CG), YF-NS3 (GCG TAA GGC TGG AAA GAG TG and CTT CCT CCC TTC ATC CAC AA), TNFA (GCC CAT GTT GTA GCA AAC CC and GGA GGT TGA CCT TGG TCT GG), IL6 (TGC AAT AAC CAC CCC TGA CC and GTG CCC ATG CTA CAT TTG CC), IFNB (AAG CAA TTG TCC AGT CCC A and TGC ATT ACC TGA AGG CCA AG), PKR (CAG AAT TGA CGG AAA GAC TTA CG and CTC TCA AGA GAA TCA TCA CTG GT), G3BP1 (CAG TTA TAT AGA CCG GCG GC and TAT GTC CAA ACC TAC GCG GC), G3BP2 (CAA TCT CAG CCA CCT CGT GT and CCA CAA CGT TTC CAA AAC TCA T), TIAR (CGT CTG GGT TAA CAG ATC AGC and CAT GGG CTG CAC TTT CAT GG), TIA1 (AAT CCC GTG CAA CAG CAG GA and AGG CGG TTG CAC TCC ATA AT). Viral genome equivalents concentrations (GE/ml) were determined by extrapolation from a standard curve generated from serial dilutions of the plasmid encoding the YF-R.luc2A-RP (62).

### Immunofluorescence, fluorescence *in situ* hybridization (FISH) and confocal analysis

Cells were fixed with 4% paraformaldehyde (PFA; Sigma-Aldrich) for 30 min at RT, permeabilized with 0.5% Triton X-100 in PBS and then blocked with 0,05% Tween-, 5% BSA-containing PBS before incubation with the indicated primary antibodies. Antibody labelling was revealed with Alexa Fluor^®^ 488-, 594 or 647-conjugated antibodies. Coverslips were mounted on slides using ProLong^®^ Gold Antifade Reagent with NucBlue solution (Invitrogen). Images were acquired with a Zeiss LSM 700 inverted confocal microscope. Fluorescence in situ hybridization was performed after immunostaining. The (+) RNA strands of YFV were detected following the manufacturer’s protocol (ViewRNA ISH Cells Assays) using probe sets designed by Affymetrix. The Alexa-Fluor 546-conjugated (+)-strand probe set targets a region between position nucleotides 4567 and 5539 of the YFV genome. Intensity plot data were generated using Zen 2 lite (Zeiss) and plotted using GraphPad Prism 6.

### Stress granule quantification

Nucleus and stress granule quantifications were performed using CellProfiler (v3.0.0) using custom pipeline (available on request). In a nutshell: (i) nuclei were detected using the NucBlue signal, (ii) cellular border were estimated using TIAR or eiF3b signal, (iii) stress granule borders were estimated using G3BP, TIAR, eIF4G or eIF3b signals. Further analyses were performed using R and R studio (63). Cells were considered positive for stress granules if they exhibited at least two of them.

### Statistical analysis

Data are presented as means ± SD and were analyzed using GraphPad Prism 6. Statistical analysis of percentage values or fold enrichment values were performed on logit or log-transformed values, respectively. Statistical analysis was performed with two tailed unpaired t-test with Welch correction for pairwise comparisons or by one- or two-way analysis of variance (ANOVA) with Tukey’s or Dunnet’s multiple comparisons test when comparing more than two sets of values. Each experiment was performed at least twice, unless otherwise stated. Statistically significant differences are indicated as follows: *: p < 0.05, **: p < 0.01 and ***: p < 0.001; ns, not significant.

## Acknowledgments

We thank Charlie Rice for generously providing the anti-YFV-NS4b antibodies; Richard Kuhn for the YFV replicon construct, Marie Flamand for anti-NS1 antibodies (17A12), Christine Neuveut for the HepG2 cells, Olivier Schwartz for the CMMT cells, Annette Martin for Huh7 cells, Pamela Schnupf for eIF3b and eIF4G antibodies, Pierre Charneau for the plasmids pCMV-VSV-G and pCMVR8.74, Andrea Cimarelli for the pALPS plasmid, Olivier Disson for immunohistochemistry advice and Léa Richard for cloning shPKR into pALPS. We are grateful to the members of our laboratory for helpful discussions and technical advice. We thank Audrey Salles and Julien Fernandes (UTechS Photonic Bioimaging platform, Institut Pasteur) for their help in confocal acquisition. We are grateful to Stéphane Rigaud (Image Analysis Hub, Institut Pasteur) for his advice in analyzing confocal images and to Vincent Guillemot (Bioinformatics and Biostatistics HUB, Institut Pasteur) for his help in statistical analysis. Finally, we thank Stephanie Straubel for critical reading of the manuscript.

## References

1. Gould EA, Solomon T. 2008. Pathogenic flaviviruses. Lancet 371:500–9.

2. Gubler DJ, Kuno G, Markoff L. 2007. Flaviviruses, p. 1153–1252. In Fields Virology 5th Edition.

3. Douam F, Ploss A. 2018. Yellow Fever Virus: Knowledge Gaps Impeding the Fight Against an Old Foe. Trends Microbiol 26:913–928.

4. Pulendran B. 2009. Learning immunology from the yellow fever vaccine: innate immunity to systems vaccinology. Nat Rev Immunol 9:741–7.

5. Barrett AD. 2016. Yellow Fever in Angola and Beyond--The Problem of Vaccine Supply and Demand. N Engl J Med 375:301–3.

6. Barrett ADT. 2018. The reemergence of yellow fever. Science 361:847–848.

7. Messaoudi I, Basler CF. 2015. Immunological features underlying viral hemorrhagic fevers. Curr Opin Immunol 36:38–46.

8. Paessler S, Walker DH. 2013. Pathogenesis of the viral hemorrhagic fevers. Annu Rev Pathol 8:411–40.

9. Chen IY, Ichinohe T. 2015. Response of host inflammasomes to viral infection. Trends Microbiol 23:55–63.

10. ter Meulen J, Sakho M, Koulemou K, Magassouba N, Bah A, Preiser W, Daffis S, Klewitz C, Bae HG, Niedrig M, Zeller H, Heinzel-Gutenbrunner M, Koivogui L, Kaufmann A. 2004. Activation of the cytokine network and unfavorable outcome in patients with yellow fever. The Journal of infectious diseases 190:1821–7.

11. Engelmann F, Josset L, Girke T, Park B, Barron A, Dewane J, Hammarlund E, Lewis A, Axthelm MK, Slifka MK, Messaoudi I. 2014. Pathophysiologic and transcriptomic analyses of viscerotropic yellow fever in a rhesus macaque model. PLoS Negl Trop Dis 8:e3295.

12. Streicher F, Jouvenet N. 2019. Stimulation of Innate Immunity by Host and Viral RNAs. Trends in Immunology 40:1134–1148.

13. Chow KT, Gale M, Loo Y-M. 2018. RIG-I and Other RNA Sensors in Antiviral Immunity. Annual Review of Immunology 36:667–694.

14. Nan Y, Nan G, Zhang YJ. 2014. Interferon induction by RNA viruses and antagonism by viral pathogens. Viruses 6:4999–5027.

15. Chazal M, Beauclair G, Gracias S, Najburg V, Simon-Loriere E, Tangy F, Komarova AV, Jouvenet N. 2018. RIG-I Recognizes the 5’ Region of Dengue and Zika Virus Genomes. Cell Rep 24:320–328.

16. Sprokholt JK, Kaptein TM, van Hamme JL, Overmars RJ, Gringhuis SI, Geijtenbeek TBH. 2017. RIG-I-like Receptor Triggering by Dengue Virus Drives Dendritic Cell Immune Activation and TH1 Differentiation. J Immunol 198:4764–4771.

17. Sprokholt JK, Kaptein TM, van Hamme JL, Overmars RJ, Gringhuis SI, Geijtenbeek TBH. 2017. RIG-I-like receptor activation by dengue virus drives follicular T helper cell formation and antibody production. PLoS Pathog 13:e1006738.

18. Hertzog J, Junior AGD, Rigby RE, Donald CL, Mayer A, Sezgin E, Song C, Jin B, Hublitz P, Eggeling C, Kohl A, Rehwinkel J. 2018. Infection with a Brazilian isolate of Zika virus generates RIG-I stimulatory RNA and the viral NS5 protein blocks type I IFN induction and signaling. European Journal of Immunology 48:1120–1136.

19. Bruni D, Chazal M, Sinigaglia L, Chauveau L, Schwartz O, Despres P, Jouvenet N. 2015. Viral entry route determines how human plasmacytoid dendritic cells produce type I interferons. Sci Signal 8:ra25.

20. Onomoto K, Jogi M, Yoo J-S, Narita R, Morimoto S, Takemura A, Sambhara S,Kawaguchi A, Osari S, Nagata K, Matsumiya T, Namiki H, Yoneyama M, Fujita T. 2012. Critical Role of an Antiviral Stress Granule Containing RIG-I and PKR in Viral Detection and Innate Immunity. PLOS ONE 7:e43031.

21. Oh S-W, Onomoto K, Wakimoto M, Onoguchi K, Ishidate F, Fujiwara T, Yoneyama M, Kato H, Fujita T. 2016. Leader-Containing Uncapped Viral Transcript Activates RIG-I in Antiviral Stress Granules. PLOS Pathogens 12:e1005444.

22. Yoo J-S, Takahasi K, Ng CS, Ouda R, Onomoto K, Yoneyama M, Lai JC, Lattmann S, Nagamine Y, Matsui T, Iwabuchi K, Kato H, Fujita T. 2014. DHX36 enhances RIG-I signaling by facilitating PKR-mediated antiviral stress granule formation. PLoS Pathog 10:e1004012.

23. Sánchez-Aparicio MT, Ayllón J, Leo-Macias A, Wolff T, García-Sastre A. 2017. Subcellular Localizations of RIG-I, TRIM25, and MAVS Complexes. J Virol 91.

24. Langereis MA, Feng Q, Kuppeveld FJ van. 2013. MDA5 Localizes to Stress Granules, but This Localization Is Not Required for the Induction of Type I Interferon. Journal of Virology 87:6314–6325.

25. Narita R, Takahasi K, Murakami E, Hirano E, Yamamoto SP, Yoneyama M, Kato H, Fujita T. 2014. A Novel Function of Human Pumilio Proteins in Cytoplasmic Sensing of Viral Infection. PLOS Pathogens 10:e1004417.

26. Hu S, Sun H, Yin L, Li J, Mei S, Xu F, Wu C, Liu X, Zhao F, Zhang D, Huang Y, Ren L, Cen S, Wang J, Liang C, Guo F. 2019. PKR-dependent cytosolic cGAS foci are necessary for intracellular DNA sensing. Sci Signal 12.

27. Onomoto K, Yoneyama M, Fung G, Kato H, Fujita T. 2014. Antiviral innate immunity and stress granule responses. Trends Immunol 35:420–428.

28. McCormick C, Khaperskyy DA. 2017. Translation inhibition and stress granules in the antiviral immune response. Nat Rev Immunol 17:647–660.

29. Ivanov P, Kedersha N, Anderson P. 2019. Stress Granules and Processing Bodies in Translational Control. Cold Spring Harb Perspect Biol 11:a032813.

30. Reineke LC, Lloyd RE. 2013. Diversion of stress granules and P-bodies during viral infection. Virology 436:255–267.

31. Sumpter R, Loo YM, Foy E, Li K, Yoneyama M, Fujita T, Lemon SM, Gale M. 2005. Regulating intracellular antiviral defense and permissiveness to hepatitis C virus RNA replication through a cellular RNA helicase, RIG-I. J Virol 79:2689–99.

32. Kedersha N, Cho MR, Li W, Yacono PW, Chen S, Gilks N, Golan DE, Anderson P. 2000. Dynamic Shuttling of Tia-1 Accompanies the Recruitment of mRNA to Mammalian Stress Granules. J Cell Biol 151:1257–1268.

33. Bonenfant G, Williams N, Netzband R, Schwarz MC, Evans MJ, Pager CT. 2019. Zika Virus Subverts Stress Granules To Promote and Restrict Viral Gene Expression. Journal of Virology 93.

34. Emara MM, Brinton MA. 2007. Interaction of TIA-1/TIAR with West Nile and dengue virus products in infected cells interferes with stress granule formation and processing body assembly. Proceedings of the National Academy of Sciences 104:9041–9046.

35. Monath TP. 2005. Yellow fever vaccine. Expert review of vaccines 4:553–74.

36. Hou S, Kumar A, Xu Z, Airo AM, Stryapunina I, Wong CP, Branton W, Tchesnokov E, Götte M, Power C, Hobman TC. 2017. Zika Virus Hijacks Stress Granule Proteins and Modulates the Host Stress Response. Journal of Virology 91.

37. Tu Y-C, Yu C-Y, Liang J-J, Lin E, Liao C-L, Lin Y-L. 2012. Blocking Double-Stranded RNA-Activated Protein Kinase PKR by Japanese Encephalitis Virus Nonstructural Protein 2A. Journal of Virology 86:10347–10358.

38. Samuel MA, Whitby K, Keller BC, Marri A, Barchet W, Williams BRG, Silverman RH, Gale M, Diamond MS. 2006. PKR and RNase L Contribute to Protection against Lethal West Nile Virus Infection by Controlling Early Viral Spread in the Periphery and Replication in Neurons. J Virol 80:7009–7019.

39. Dabo S, Meurs EF. 2012. dsRNA-Dependent Protein Kinase PKR and its Role in Stress, Signaling and HCV Infection. Viruses 4:2598–2635.

40. Lu L, Han AP, Chen JJ. 2001. Translation initiation control by heme-regulated eukaryotic initiation factor 2alpha kinase in erythroid cells under cytoplasmic stresses. Mol Cell Biol 21:7971–7980.

41. Bonnet MC, Weil R, Dam E, Hovanessian AG, Meurs EF. 2000. PKR stimulates NF-kappaB irrespective of its kinase function by interacting with the IkappaB kinase complex. Mol Cell Biol 20:4532–4542.

42. Bonnet MC, Daurat C, Ottone C, Meurs EF. 2006. The N-terminus of PKR is responsible for the activation of the NF-ĸB signaling pathway by interacting with the IKK complex. Cellular Signalling 18:1865–1875.

43. Sinigaglia L, Gracias S, Decembre E, Fritz M, Bruni D, Smith N, Herbeuval JP, Martin A, Dreux M, Tangy F, Jouvenet N. 2018. Immature particles and capsid-free viral RNA produced by Yellow fever virus-infected cells stimulate plasmacytoid dendritic cells to secrete interferons. Sci Rep 8:10889.

44. Errett JS, Suthar MS, McMillan A, Diamond MS, Gale M. 2013. The essential, nonredundant roles of RIG-I and MDA5 in detecting and controlling West Nile virus infection. J Virol 87:11416–25.

45. Zamanian-Daryoush M, Mogensen TH, DiDonato JA, Williams BR. 2000. NF-kappaB activation by double-stranded-RNA-activated protein kinase (PKR) is mediated through NF-kappaB-inducing kinase and IkappaB kinase. Mol Cell Biol 20:1278–1290.

46. Pham AM, Santa Maria FG, Lahiri T, Friedman E, Marié IJ, Levy DE. 2016. PKR Transduces MDA5-Dependent Signals for Type I IFN Induction. PLoS Pathog 12.

47. Gil J, García MA, Gomez-Puertas P, Guerra S, Rullas J, Nakano H, Alcamí J, Esteban M. 2004. TRAF Family Proteins Link PKR with NF-ĸB Activation. Molecular and Cellular Biology 24:4502–4512.

48. Arnaud N, Dabo S, Akazawa D, Fukasawa M, Shinkai-Ouchi F, Hugon J, Wakita T, Meurs EF. 2011. Hepatitis C virus reveals a novel early control in acute immune response. PLoS Pathog 7:e1002289.

49. Zhang P, Li Y, Xia J, He J, Pu J, Xie J, Wu S, Feng L, Huang X, Zhang P. 2014. IPS-1 plays an essential role in dsRNA-induced stress granule formation by interacting with PKR and promoting its activation. J Cell Sci 127:2471–2482.

50. Ravindran R, Khan N, Nakaya HI, Li S, Loebbermann J, Maddur MS, Park Y, Jones DP, Chappert P, Davoust J, Weiss DS, Virgin HW, Ron D, Pulendran B. 2014. Vaccine activation of the nutrient sensor GCN2 in dendritic cells enhances antigen presentation. Science 343:313–317.

51. Albornoz A, Carletti T, Corazza G, Marcello A. 2014. The stress granule component TIA-1 binds tick-borne encephalitis virus RNA and is recruited to perinuclear sites of viral replication to inhibit viral translation. J Virol 88:6611–6622.

52. Roth H, Magg V, Uch F, Mutz P, Klein P, Haneke K, Lohmann V, Bartenschlager R, Fackler OT, Locker N, Stoecklin G, Ruggieri A. 2017. Flavivirus Infection Uncouples Translation Suppression from Cellular Stress Responses. mBio 8.

53. Basu M, Courtney SC, Brinton MA. 2017. Arsenite-induced stress granule formation is inhibited by elevated levels of reduced glutathione in West Nile virus-infected cells. PLOS Pathogens 13:e1006240.

54. Amorim R, Temzi A, Griffin BD, Mouland AJ. 2017. Zika virus inhibits eIF2a-dependent stress granule assembly. PLOS Neglected Tropical Diseases 11:e0005775.

55. Zhang Q, Sharma NR, Zheng Z-M, Chen M. 2019. Viral Regulation of RNA Granules in Infected Cells. Virol Sin 34:175–191.

56. Christ W, Tynell J, Klingström J. 2020. Puumala and Andes Orthohantaviruses Cause Transient Protein Kinase R-Dependent Formation of Stress Granules. J Virol 94.

57. Blight KJ, McKeating JA, Rice CM. 2002. Highly permissive cell lines for subgenomic and genomic hepatitis C virus RNA replication. J Virol 76:13001–14.

58. Pertel T, Hausmann S, Morger D, Zuger S, Guerra J, Lascano J, Reinhard C, Santoni FA, Uchil PD, Chatel L, Bisiaux A, Albert ML, Strambio-De-Castillia C, Mothes W, Pizzato M, Grutter MG, Luban J. 2011. TRIM5 is an innate immune sensor for the retrovirus capsid lattice. Nature 472:361–5.

59. Jones CT, Patkar CG, Kuhn RJ. 2005. Construction and applications of yellow fever virus replicons. Virology 331:247–59.

60. RStudio Team. 2016. RStudio: Integrated Development Environment for R.

